# Intravenous administration of human umbilical cord mesenchymal stromal cells leads to an inflammatory response in the lung

**DOI:** 10.1101/2022.09.26.509547

**Authors:** Alejandra Hernandez Pichardo, Bettina Wilm, Neill Liptrott, Patricia Murray

## Abstract

Mesenchymal stromal cells (MSCs) administered intravenously (IV) have shown efficacy in pre-clinical models of various diseases. This is despite the cells not reaching the site of injury due to entrapment in the lungs. The ability of MSCs to modulate immune responses has been proposed as one of the mechanisms by which these cells provide therapeutic benefits, irrespective of whether they are sourced from bone marrow, adipose tissue or umbilical cord. To better understand how MSCs affect innate immune cell populations in the lung, we evaluated the percentage, distribution and phenotype of neutrophils, monocytes and macrophages by flow cytometry and histological analyses after delivering human umbilical cord-derived MSCs (hUC-MSCs) IV into immunocompetent mice. After 2 h, we observed a sharp increase in neutrophils, and pro-inflammatory monocytes and macrophages. Moreover, these immune cells localised in the vicinity of the MSCs suggesting an active role in their clearance. By 24 h, we detected an increase in anti-inflammatory monocytes and macrophages. These results suggest that the IV injection of hUC-MSCs leads to an initial inflammatory phase in the lung shortly after injection, followed by a resolution phase 24 h later.

## 1. Introduction

After intravenous (IV) injection, most non-haematological cells, including mesenchymal stromal cells (MSCs) remain trapped within the lung vasculature and die within 24 h (1,2). This is referred to as the first-pass effect (3), which disproves the theory that MSCs migrate to the site of injury and differentiate into local cells resulting in a regenerative response (4). Evidence suggest that the therapeutic benefits of MSCs is likely mediated, at least in part, by the release of trophic factors and their ability to modulate the immune system (5–7).

Several studies suggest that exogenous MSCs can ameliorate injury in a variety of animal models such as the heart (8), eye (9), kidney (10), bone (11), cartilage (12), liver (13), amongst others. The mechanisms are not fully elucidated and understanding the initial effect that MSCs have on the immune-cell populations in the lung after IV delivery could shed light on this question.

To provide an early line of defence against infections in tissues, the first type of immune cells that respond to invading pathogens or foreign materials are the cells of the innate immune system (14). Bone marrow-derived myeloid cells consist of a heterogeneous population comprising monocytes, macrophages and dendritic cells (DCs), as well as granulocytes (mast cells, basophils, eosinophils, and neutrophils) (15). Despite each subset having specialized functions based on their environment, all myeloid cells play a role in the phagocytosis of foreign materials, opsonised extracellular microbes, and dying/dead cells (16–18). Moreover, they secrete cytokines and chemokines to induce an immune response (19).

Neutrophils, the most abundant type of granulocytes, are the first cells recruited to sites of injury, followed by monocytes and macrophages (20,21). Neutrophils are known for their microbe-clearing mechanisms involving reactive oxygen species generation, antimicrobial protein degranulation, and neutrophil extracellular traps (NETs) formation (22,23). MSCs have been shown to mediate their therapeutic benefits by modulating neutrophils; for instance, bacterial clearance was enhanced in a murine sepsis model after MSC IV administration because the MSCs enhanced the phagocytic capacity of the neutrophils (24).

Monocytes are precursor cells that give rise to DCs and macrophages. Their mobility gives them a unique role in the mononuclear phagocyte system. In contrast to the limited migration potential of terminally differentiated DCs and macrophages, monocytes are rapidly mobilized upon challenge and can access any location within the body catering to its needs (25). In the context of cell therapies, an IV injection of hUC-MSCs into mice showed that monocytes mediate the rapid clearance of the cells by phagocytosis (26). Moreover, phagocytosis of the administered cells in the lung resulted in monocyte reprogramming toward an anti-inflammatory activation state (26).

Tissue macrophages play various homeostatic roles such as tissue remodelling and repair, clearing of senescent cells, as well as induction and resolution of the inflammatory response (25). In addition, macrophages engage with T and B lymphocytes and participate in the induction of adaptive immunity (27). The IV infusion of MSCs in mice leads to an inflammatory response accompanied by increased numbers of macrophages in the lungs shortly after the cells home to this organ (28). Moreover, at 1 week following administration the MSCs increase the number of anti-inflammatory macrophages and decrease the inflammatory macrophages *in vivo* (29), affecting disease outcomes via macrophage polarization (30). Also, MSCs alleviated tissue damage and inflammation in a mouse model of acute kidney injury by inducing macrophage polarization toward an anti-inflammatory phenotype within 24 h following IV administration of the MSCs (31).

Macrophages can further be categorized based on their location. In the lung, resident alveolar macrophages are maintained by local proliferation (32) and perform tissue-specific roles such as surfactant clearance (33). MSCs reduced the severity of lung injury in an *E. coli* pneumonia model and modulated alveolar macrophage polarization *in vivo* (34). Interstitial macrophages mature in the lung after the recruitment of precursors from the blood (25). Their localization within the lung remains unclear but studies have shown their presence in the parenchyma (35) and the bronchial interstitium (36). Their functions include phagocytosis of foreign invaders, antigen presentation and immune modulation (37).

Some studies have investigated the interactions between infused MSCs and specific immune cell populations *in vivo* (26,38,39) but a comprehensive analysis on the impact on innate cells is lacking, especially in the period immediately following MSC administration. In this study, we have used hUC-MSCs because of reports suggesting that they have superior immunomodulatory properties than those derived from bone marrow (40). Our aim was to investigate the fate of hUC-MSCs in the lung and their effect on the proportion, distribution and polarization state of innate immune cell populations, particularly granulocytes, monocytes and macrophages. To do this, we administered cells into immunocompetent naïve mice. Then, we measured the changes in myeloid cells at two different time points: 2 hours and 24 hours. These time points were selected because at 2 hours following cell injection, most cells remain viable, while by 24 hours, most of the hUC-MSCs have been cleared.

## 2. Methods

### 2.1 Cell preparation

Primary human umbilical cord-derived mesenchymal stromal cells (hUC-MSCs) were collected from consenting donors at the National Health Service Blood and Transplant (NHSBT, UK) and obtained at passage 3.

The hUC-MSCs were transduced in the presence of 6 μg/ml DEAE-Dextran with a lentiviral vector pCDH-EF1-Luc2-P2A-tdTomato, encoding luc2 firefly luciferase (FLuc) reporter under the constitutive elongation factor 1-α (EF1α) promoter and upstream of a P2A linker followed by the tdTomato fluorescent protein (gift from Kazuhiro Oka;Addgene plasmid #72486; http://n2t.net/addgene:72486;RRID:Addgene_72486). To obtain a >98% transduced population, the cells were sorted based on tdTomato fluorescence (BD FACS Aria). The cells were cultured in α-MEM supplemented with 10% FBS and incubated at 37°C, 5% CO2. When confluent, cells were washed with PBS and incubated with 1% trypsin at 37°C for 3 min. Subsequently, culture medium was added and the cell suspension was transferred to a 15 ml conical tube and centrifuged at 300G for 3 minutes. The supernatant was discarded and the cell pellet resuspended in an adequate volume of medium for passaging. Prior to re-plating, 10 μl of cell suspension was transferred into a haemocytometer and cells were counted. All procedures were performed under sterile conditions.

### 2.2 Animal studies

All experiments were carried out under a licence granted under the UK Animals Act 1986 and were approved by the ethics committee of the University of Liverpool Animal Welfare and Ethics Review Board (AWERB). Eight to ten-week-old female albino mice (C57BL/6) (B6N-TyrC-Brd/BrdCrCrl, originally purchased from the Jackson Lab) were housed in individually ventilated cages under a 12 h light/dark cycle, with ad libitum access to water and food.

### 2.3 Dissociation of lung tissue

Mice received an IV injection of 2.5 × 10^5^ untransduced hUC-MSCs or PBS (100 μL) under anaesthesia with isoflurane. 2 or 24 h post injection, the animals were culled by cervical dislocation. The lungs were removed *en bloc*. The large airways were dissected from the peripheral lung tissue and each lung lobe was separated. The lung lobes were cut into small pieces with scissors, transferred into C-tubes (Miltenyi Biotec), and processed in digestion buffer (1 mg/ml of Collagenase D and 80 U/ml DNase I, both from Roche, in DMEM) and a GentleMACS dissociator (Miltenyi Biotec), according to the manufacturer’s instructions. The lung homogenates were strained through a 70 mm nylon mesh to obtain single-cell suspensions. Red blood cells were lysed using ammonium-chloride-potassium (ACK) lysis buffer (Gibco, A1049201). The resultant cells were counted using an automated cell counter (TC10, BioRad).

### 2.4 Flow cytometry

One million cells suspended in 90 μl of staining buffer (eBiosciences, 00-4222-26) were incubated with 10 μl FcBlock (Miltenyi Biotec, 130-092-575) to reduce nonspecific antibody binding. The cells were stained with a mixture of fluorochrome-conjugated antibodies (see Table 1 for a list of antibodies, clones, and fluorochromes). Data were acquired on a BD CANTO II flow cytometer using BD FACSDiva software (BD Biosciences; see Supplementary figure 1 for instrument configuration), and compensation and data analyses were performed using the DIVA software). The gating strategy followed was adapted from Misharin, A. *et al*. 2013 (42). The gating strategy used to identify cell populations is shown in Figure 2 and Figure *3*.

**Table 1.**
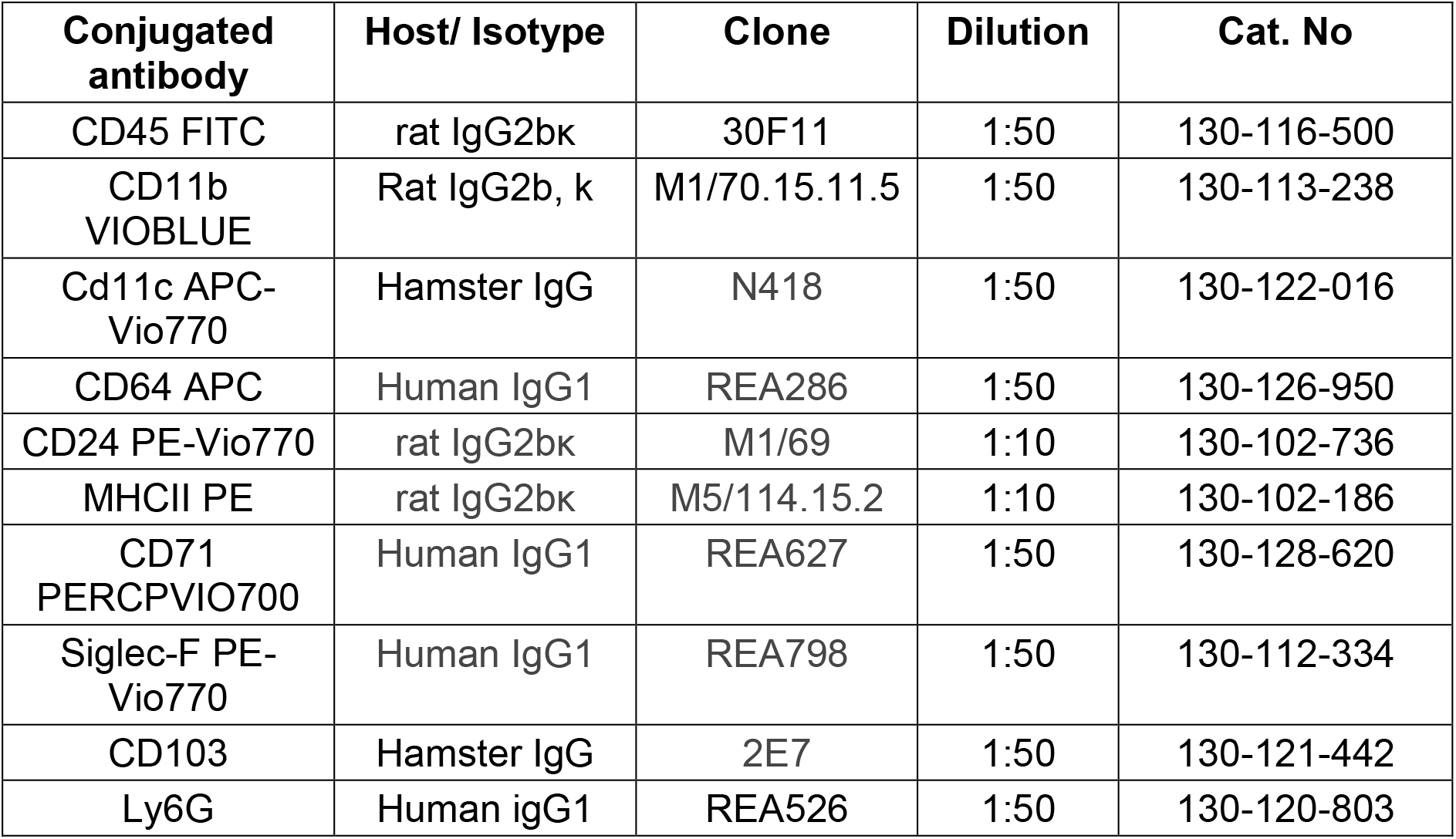
Antibodies used for flow cytometry. All antibodies were purchased from Miltenyi Biotec

| **Conjugated antibody** | **Host/ Isotype** | **Clone** | **Dilution** | **Cat. No** |
| --- | --- | --- | --- | --- |
| CD45 FITC | rat IgG2bk | 30F11 | 1:50 | 130-116-500 |
| CD11b<br>VIOBLUE | Rat IgG2b, k | M1/70.15.11.5 | 1:50 | 130-113-238 |
| Cd11c APC-Vio770 | Hamster IgG | N418 | 1:50 | 130-122-016 |
| CD64 APC | Human IgG1 | REA286 | 1:50 | 130-126-950 |
| CD24 PE-Vio770 | rat IgG2bk | M1/69 | 1:10 | 130-102-736 |
| MHCII PE | rat IgG2bk | M5/114.15.2 | 1:10 | 130-102-186 |
| CD71<br>PERCPVIO700 | Human IgG1 | REA627 | 1:50 | 130-128-620 |
| Siglec-F PE-Vio770 | Human IgG1 | REA798 | 1:50 | 130-112-334 |
| CD103 | Hamster IgG | 2E7 | 1:50 | 130-121-442 |
| Ly6G | Human igG1 | REA526 | 1:50 | 130-120-803 |

### 2.5 Retrograde perfusion fixation

2 or 24 h after IV administration of FLuc+ tdTomato hUC-MSCs, the mice received an intraperitoneal overdose of pentobarbital (Pentoject, 100 μl) followed by cannulation of the abdominal aorta, snipping of the vena cava and flushing of Heparin/PBS (5 IU/ml) with a manual pump at a constant pressure of 200 mbar (Supplementary figure 2) for 6 min to remove all blood cells, followed by 6 min perfusion with 4% w/v paraformaldehyde (PFA) to fix the whole animal. The total volume of each solution used per animal was 40 ml. The trachea was tied tightly with a surgical suture before opening the thoracic cavity for lung dissection. Finally, the lungs were post-fixed in 4% PFA overnight at 4°C.

### 2.6 Immunofluorescence

Before staining, the lungs were cleared using the CUBIC protocol (43). The cleared lungs were sucrose protected and cryo-embedded in optimal cutting temperature (OCT) medium before sectioning 30 μm thick sections using a cryostat (Thermo Scientific, Microm HM505E) at −20°C and stored at −80°C.

All sections were washed with PBS 3x for 5 min. Tissues were incubated with primary antibodies for 2 h at RT or O/N at 4°C. The primary antibody was washed with PBS, and the secondary antibodies and DAPI were added. Incubation was done at RT for 1 h. After a final washing step, the sections were mounted in fluorescence mounting media (Dako, S3023). All antibodies and dilutions used can be found in Table 2.

**Table 2.** Antibodies used for immunofluorescence.

|  | **Host** | **Clone** | **Isotype** | **Dilution** | **T°/time** | **Manufacturer** |
| --- | --- | --- | --- | --- | --- | --- |
| F4/80 - FITC | Human | REA126 | igG1 | 1:50 | RT / 2 h | Miltenyi biotech (130-117-509) |
| CD16/32 | Human | REA377 | igG1 | 1:10 | RT / 2 h | Miltenyi biotech (130-107-066) |
| Ly6C | Rat | HK1.4 | IgG2c, k | 1:200 | 4 / ON | Biolegend (128001) |
| Myeloperoxidase (MPO) | Goat | Polyclonal | igG | 1:100 | 4 / ON | R&D Systems (AF3667-SP) |
| HDAC2 | Rabbit | Polyclonal | igG | 1:250 | RT / 2 h | Sigma-Aldrich (HPA011727) |
| CD11b - APC | Rat | M1/70.15.11.5 | IgG2b, k | 1:50 | RT / 2 h | Miltenyi biotech (130-113-231) |
| Ly6G - APC | Human | REA526 | igG1 | 1:50 | RT / 2 h | Miltenyi biotech (130-120-803) |
| CD163 | Rabbit | Polyclonal | igG | 1:100 | RT / 2 h | Invitrogen (PA5-78961) |
| Alexa Fluor® 750 donkey anti-rat | Donkey | Polyclonal | IgG | 1:200 | RT / 1 h | Abcam (ab175750) |
| Alexa Fluor® 647 goat anti-rabbit | Goat | Polyclonal | igG | 1:1000 | RT / 1 h | Invitrogen (A-212465) |
| Alexa Fluor® 647 goat anti-hamster | Goat | Polyclonal | igG | 1:1000 | RT / 1 h | Invitrogen (A-21451) |
| Alexa Fluor® 633 goat anti-rat | Goat | Polyclonal | igG | 1:1000 | RT / 1 h | Invitrogen (A-21094) |

### 2.7 Imaging

Confocal microscopy images were acquired using a Leica DMi8 with Andor Dragonfly spinning disk, coupled to an EMCCD camera using a 40x/1.3 oil objective. Z-stacks were captured using the 488, 561 and 637 nm laser lines. The emission filters used were 525/50, 600/50 and 700/75. Maximum intensity projections, three-dimensional reconstructions and image analysis were done using the IMARIS (Bitplane) software package.

Cell counting with IMARIS was performed by opening *Z*-stacks in their native format, as they are automatically reconstructed into a multi-channel 3D model, which eliminates the need for image pre-processing. To designate individual cells of interest, the Spots creation tool was used. In the Spots creation wizard, the source channel corresponding to the staining of interest was selected. Background subtraction was used to separate the cell from the background. The auto-threshold value was utilized during background subtraction. The generated spots were a direct map of the intensity distribution of the immunostaining of interest as detected by Imaris. Adjustments to the Spots to create an accurate representation of the staining were made using the manual spot creation/deletion tool.

### 2.8 Statistics

Data were analysed using GraphPad Prism for Windows version 8.4.2 (GraphPad Software, Inc., San Diego, CA). Values are presented as means ± standard deviations. Comparisons between animal groups were performed using the Kruskal-Wallis test with multiple comparisons. P < 0.05 was considered statistically significant. The number of replicates included in the analyses are given in the figure legends.

#### Data availability

The data that support the findings of this study are available to download from Zenodo at http://doi.org/10.5281/zenodo.7113094

## 3. Results

### 3.1 Effect of hUC-MSCs on the distribution of myeloid cells in the lungs

To investigate changes in biodistribution of the myeloid cells within the lung after hUC-MSC administration, the lungs of mice that received FLuc TdTomato-expressing hUC-MSCs were fixed and used to prepare frozen sections for histology.

Using the CD11b pan-myeloid marker (44), confocal microscopy revealed that 2 h post hUC-MSC IV administration, there was an increase in myeloid cells. These cells persisted in the lungs up to 24 h. Moreover, the cells accumulated in close proximity to the hUC-MSC clusters and fragments (Figure 1a), suggesting that these cells might be phagocytosing the hUC-MSCs. Quantification of myeloid cells confirmed a sharp increase in these cells at 2 h that was sustained at 24 h (Figure 1b).

**Figure 1.**
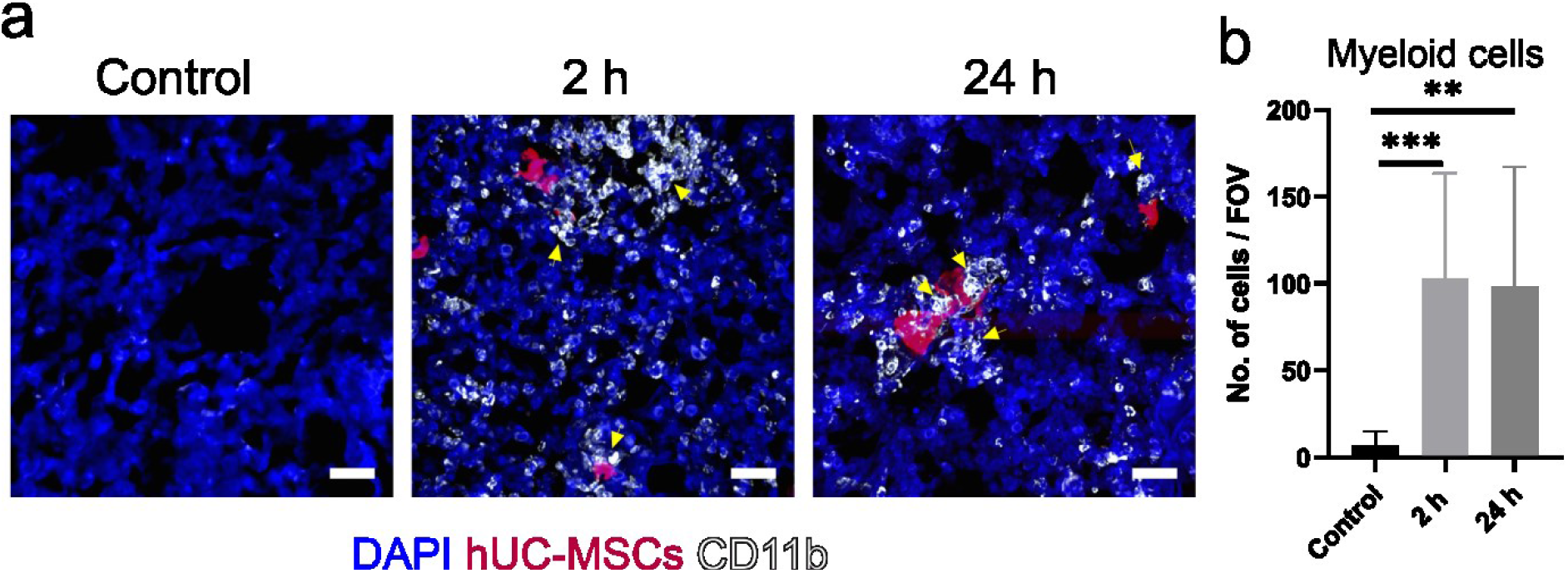
CD11b^+^ myeloid cells infiltrated into the lungs after IV administration of hUC-MSCs. a) Representative maximum intensity projection (MIP) confocal microscopy images showed CD11b^+^ cells (white) infiltrated into the lungs in comparison with control animals that received saline. CD11b^+^ cells clustered around the hUC-MSCs (red) in the lungs 2 and 24 h after cell injection (yellow arrows). Scale bar = 30 μm. b) Immunofluorescence quantification of CD11b^+^ cells. Kruskal-Wallis test with multiple comparisons. n=9; P<0.005 **; P<0.0005 ***.

Immunofluorescence analysis and quantification showed that myeloid cells infiltrate the lungs after IV injection of hUC-MSCs.

### 3.2 Implementation of gating strategy to identify myeloid cell subtypes and their polarization state

To understand what type of myeloid cells had accumulated in the lung, we performed side by side flow cytometry and immunofluorescence-based analysis of cell biodistribution within the tissue. To determine the proportion of innate immune cell sub-types, mice received untransduced hUC-MSCs IV and their lungs were dissociated for flow analysis. The markers used to identify specific populations can be found in Table 3 and the gating strategy followed is shown in figures 2 and 3. To study the cell biodistribution in the lung histologically, FLuc TdTomato-expressing hUC-MSCs were injected into a different cohort of mice.

**Table 3.**
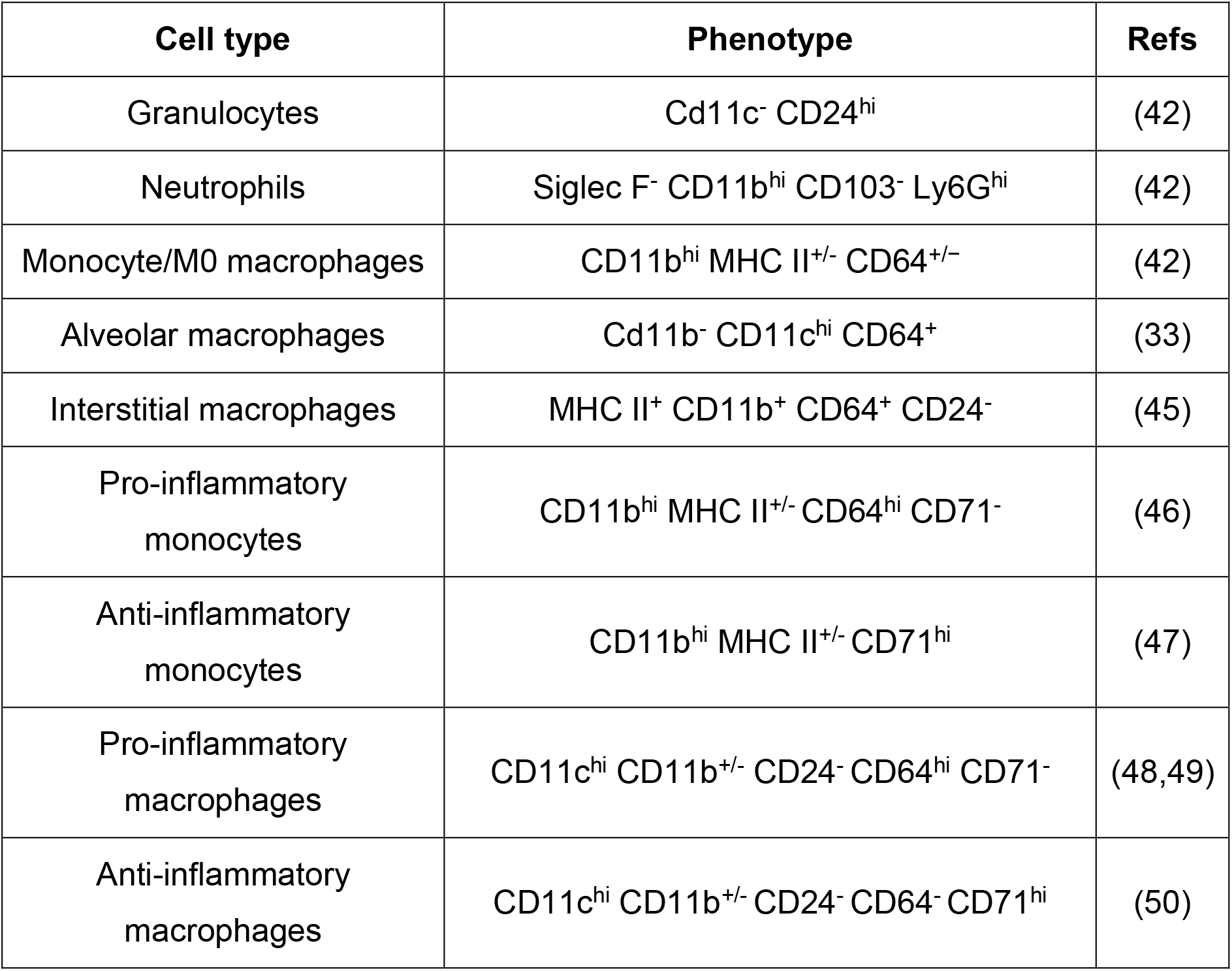
Surface markers used to identify immune cells by flow cytometry.

| **Cell type** | **Phenotype** | **Refs** |
| --- | --- | --- |
| Granulocytes | Cd11c<sup>-</sup> CD24<sup>hi</sup> | (42) |
| Neutrophils | Siglec F<sup>-</sup> CD11b<sup>hi</sup> CD103<sup>-</sup> Ly6G<sup>hi</sup> | (42) |
| Monocyte/M0 macrophages | CD11b<sup>hi</sup> MHC II<sup>+/-</sup> CD64<sup>+/-</sup> | (42) |
| Alveolar macrophages | Cd11b<sup>-</sup> CD11c<sup>hi</sup> CD64<sup>+</sup> | (33) |
| Interstitial macrophages | MHC II<sup>+</sup> CD11b<sup>+</sup> CD64<sup>+</sup> CD24<sup>-</sup> | (45) |
| Pro-inflammatory monocytes | CD11b<sup>hi</sup> MHC II<sup>+/-</sup> CD64<sup>hi</sup> CD71<sup>-</sup> | (46) |
| Anti-inflammatory monocytes | CD11b<sup>hi</sup> MHC II<sup>+/-</sup> CD71<sup>hi</sup> | (47) |
| Pro-inflammatory macrophages | CD11c<sup>hi</sup> CD11b<sup>+/-</sup> CD24<sup>-</sup> CD64<sup>hi</sup> CD71<sup>-</sup> | (48,49) |
| Anti-inflammatory macrophages | CD11c<sup>hi</sup> CD11b<sup>+/-</sup> CD24<sup>-</sup> CD64<sup>-</sup> CD71<sup>hi</sup> | (50) |

**Figure 2.**
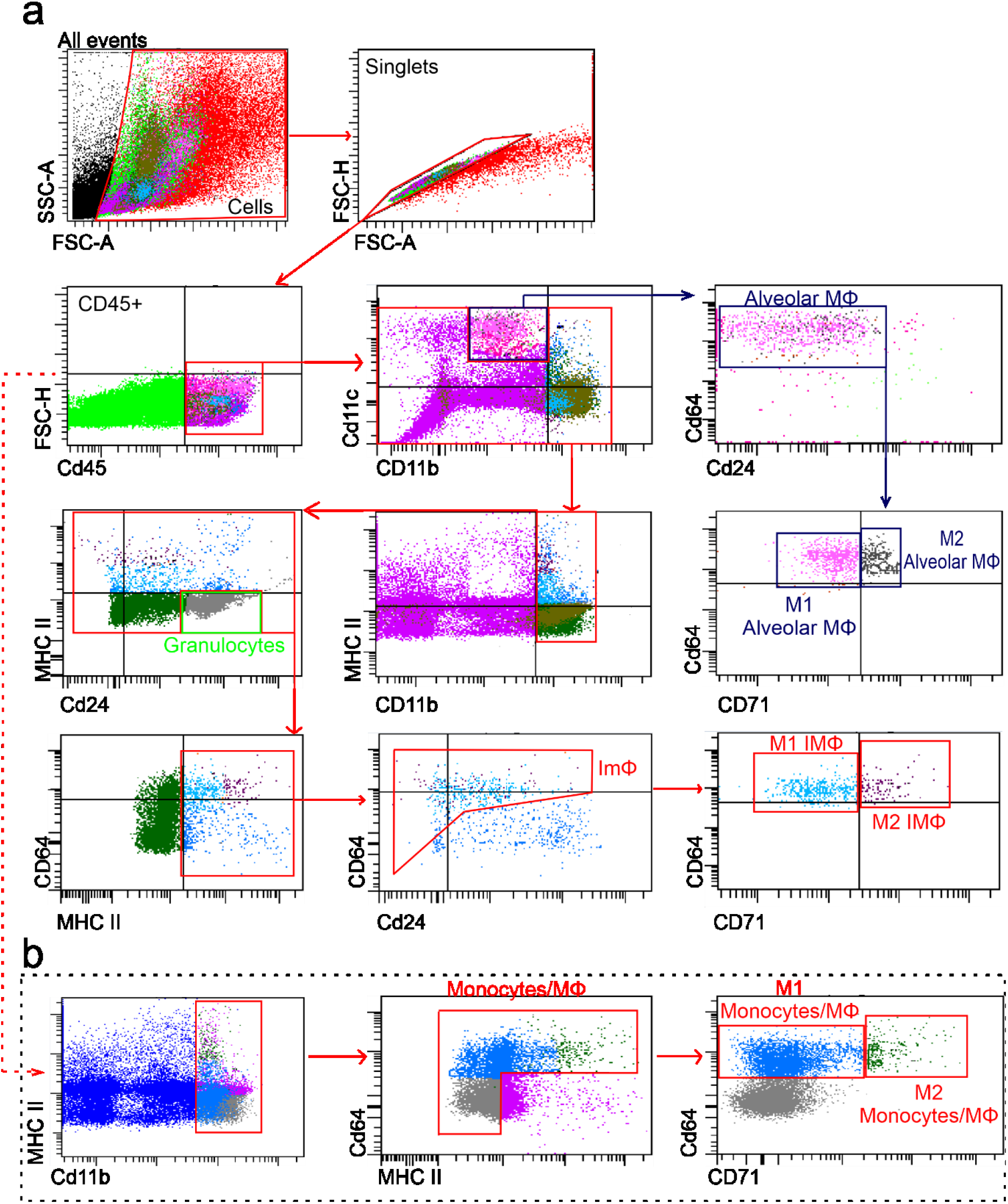
Gating strategy used to identify immune-cell subsets in the mouse lung after IV administration of hUC-MSCs or saline. After enzymatic and mechanical digestion of mouse lungs, debris and doublets were excluded. Leukocytes were identified by CD45 staining. a) To find specific populations, a sequential gating strategy was used: alveolar macrophages (MΦ) (CD11b^-^ CD11c^hi^), granulocytes (CD11c^-^ CD24^hi^), interstitial macrophages (CD11b^+^ MHC II^+^ CD64^+^ CD24^-^). The polarization status toward a pro- or anti-inflammatory phenotype was assessed by the expression of CD64 and CD71, respectively. b) A parallel gating strategy was used to identify monocytes: monocytes/M0 MΦ (CD11b^hi^ MHC II^+/-^ CD64^+/-^). To identify classically- and alternatively-activated cell types the CD64 (M1 marker) and CD71 (M2 marker) were used.

**Figure 3.**
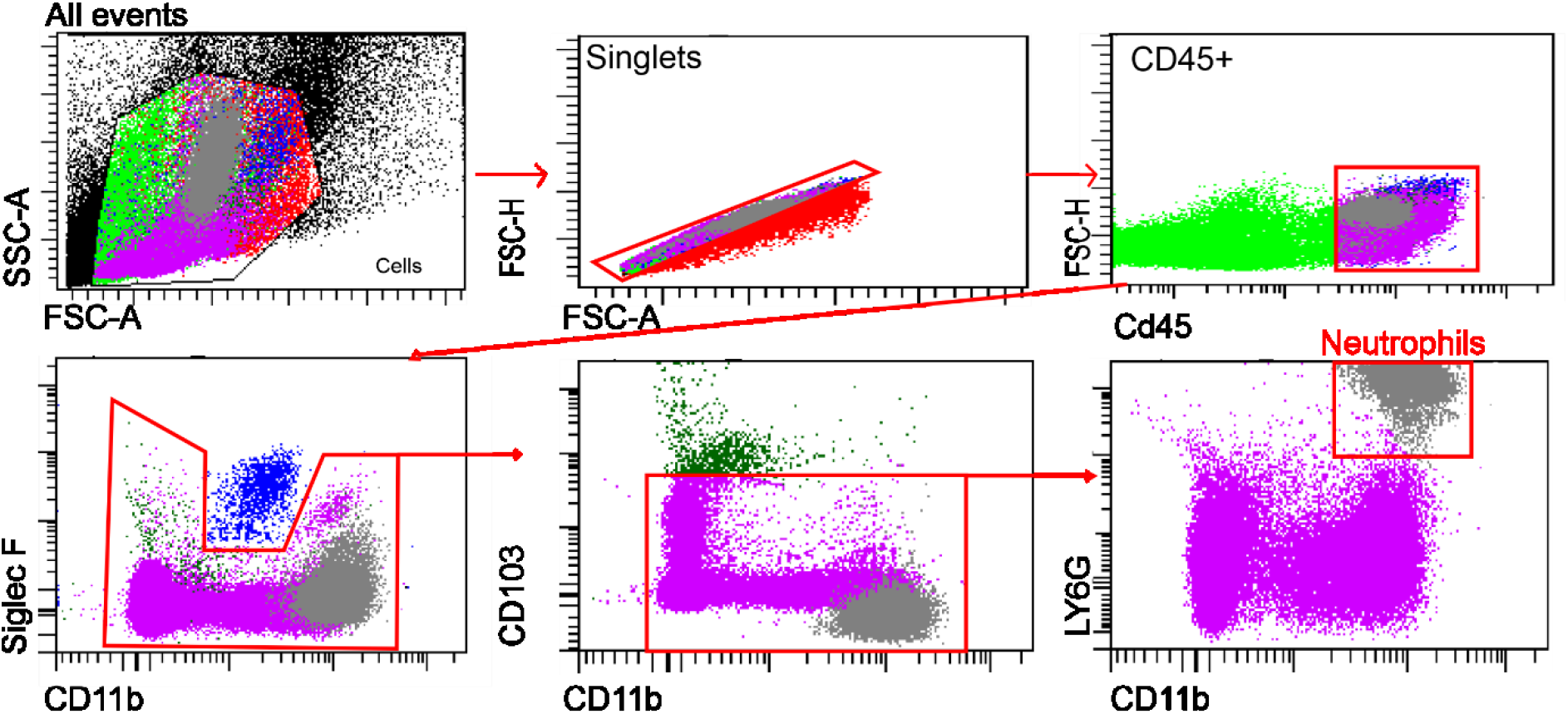
Gating strategy used to identify neutrophils in the mouse lung after IV injection of hUC-MSCs or saline. Lung enzymatic and mechanic digestion was performed. Debris and doublets were excluded. CD45 staining was used to identify leukocytes. Neutrophils were gated as Siglec F^-^ CD11b^hi^ CD103^-^ Ly6G^hi^.

The flow cytometry gating strategy used to identify myeloid cells, their specific subtypes and their polarisation states is presented here for the first time. The strategy for granulocytes, polarised resident lung macrophages and polarised interstitial macrophages is shown in Figure 2a; the strategy for polarised infiltrating monocytes/macrophages is shown in Figure 2b; and the strategy for neutrophils is shown in Figure 3.

### 3.3 Effect of hUC-MSCs on the proportion, distribution and polarization of infiltrating granulocytes within the lung

Flow cytometric analysis revealed that the proportion of granulocytes increased approximately two-fold from baseline 2 h post hUC-MSC administration. The number of these cells decreased after 24 h, but remained 1.7x higher than in controls (Figure 4a, left).

**Figure 4.**
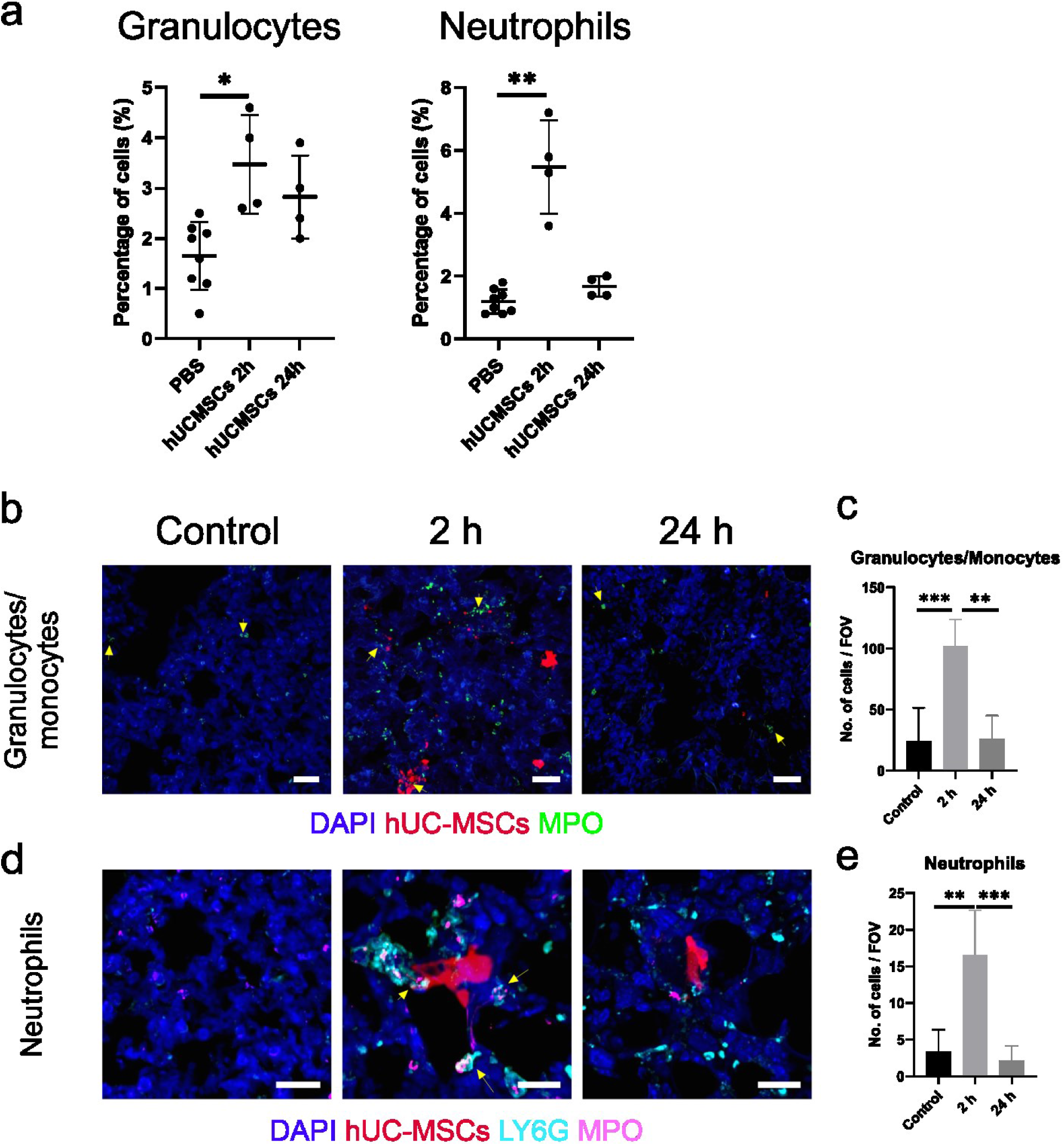
MPO positive cells are recruited to the lungs rapidly after hUC-MSC IV infusion. a) Flow cytometry showed that Cd11c^-^ CD24^hi^ granulocytes and Siglec F^-^ CD11b^hi^ CD103^-^ Ly6G^hi^ neutrophils increased 2 h after hUC-MSC injection, and decreased at the 24 h time point. Kruskal-Wallis test with multiple comparisons; control n=8, hUC-MSC group n=4. P<0.05 *; P<0.005 **. b) MPO^+^ granulocytes/monocytes (green) clustered around the hUC-MSCs (red) in the lungs 2 h after cell injection (yellow arrows) and their levels decreased at 24 h. Scale bar = 30 μm. c) Immunofluorescence quantification of MPO^+^ cells. Kruskal-Wallis test with multiple comparisons. n=9. P<0.005 **; P<0.0005 ***. d) Ly6G + MPO (magenta + cyan) neutrophils surrounded the hUC-MSCs (red) in the lungs 2 h after cell injection (yellow arrows). 24 h later, neutrophil numbers had lowered but Ly6G cells were still present in the lung. e) Immunofluorescence quantification of MPO + LY6G neutrophils. Kruskal-Wallis test with multiple comparisons. n=9. P<0.005 **; P<0.0005 ***.

The granulocyte/monocyte marker MPO (51,52) was used for immunofluorescence staining. MPO positive cells localized in the vicinity of the hUC-MSC clusters as well as in areas were cell debris was observed (Figure 4b), potentially indicating an active role of these cells in the clearance of the exogenously administered human cells. Quantification of the fluorescence images showed an approximate 3-fold increase in MPO expressing cells at 2 h, with a decline back to control levels at 24 h (Figure 4c). The higher increase observed by histology in comparison with flow cytometry can be explained as the gating strategy excluded monocytes while this was not possible using the MPO marker in immunofluorescence.

Neutrophils showed an approximate 4.5x increase at 2 h with the number of these cells returning to baseline after 24 h (Figure 4a, right). To reliably identify neutrophils by immunostaining of the lung sections, we used the MPO surface marker in combination with Ly6G, recognized as a marker that is highly expressed by neutrophils (53). We observed that double-labelled neutrophils localized to the vicinity of the hUC-MSCs at 2 h. After 24 h, cells expressing only Ly6G, which might be monocytes or other granulocytes, were observed distributed evenly throughout the lung (Figure 4d). In agreement with the flow cytometry data, their quantification confirmed that these cells increased by 4x in the lungs 2 h after cell delivery and returned to baseline levels at 24 h (Figure 4e).

Together, these data showed that granulocytes, particularly neutrophils, accumulated in the lung in response to hUC-MSC lV administration at the 2 h time point. After 24 h, the number of these cells decreased.

### 3.4 Effect of hUC-MSC on neutrophil extracellular trap formation

Given the high influx of neutrophils into the lung after IV injection of hUC-MSCs, we questioned whether NETs formed as a consequence. We stained frozen lung sections of mice that had received FLuc TdTomato-expressing hUC-MSCs for Histone deacetylase 2 (HDAC2) and MPO, common NET markers (*54*).

Although an increase in HDAC2 was observed at both time points, the characteristic elongated NET structures were not observed (*Figure 5*). Given that the HDAC2 expression colocalized with the hUC-MSCs, we speculate that as the injected cells died, their DNA was exposed leading to the observation of HDAC2 in the lung sections. Thus, NET formation was likely not induced by the administration of hUC-MSCs.

**Figure 5.**
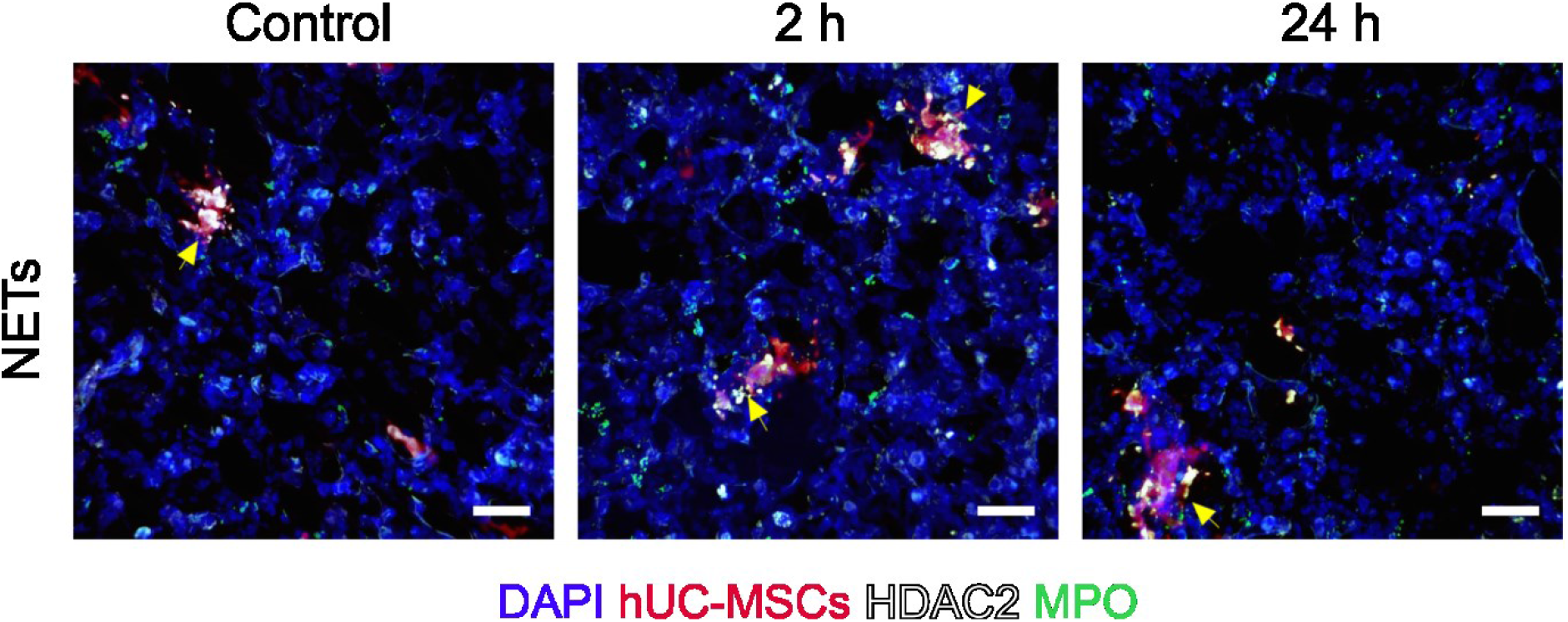
Neutrophil extracellular traps are not observed in the lungs of mice at any timepoint after hUC-MSC IV infusion. hUC-MSCs (red), MPO (green), HDAC II (white). Scale bar = 30 μm.

### 3.5 Effect of hUC-MSCs on the proportion, distribution and polarization of infiltrating monocytes and macrophages within the lung

Then, we determined how hUC-MSCs affected the quantity, localization, and phenotype of infiltrating macrophages and monocytes in the mouse lung. Monocytes and macrophages express similar surface molecules. The selection of markers used in our panel, made it difficult to differentiate between these cell populations, thus they were analysed as one population - Monocytes/M0 macrophages.

We observed by flow cytometry that 2 h after hUC-MSC administration, the proportion of monocytes/M0 macrophages increased by approximately 2.8x when compared to control. After 24 h, there was a sharp decrease (Figure 6a, left). Pro-inflammatory monocytes/M0 macrophages increased ~2-fold at 2 h (Figure 6a, middle), followed by a shift of this population toward an M2 phenotype at 24 h, with an approximate 3.2x increase (Figure 6a, right).

**Figure 6.**
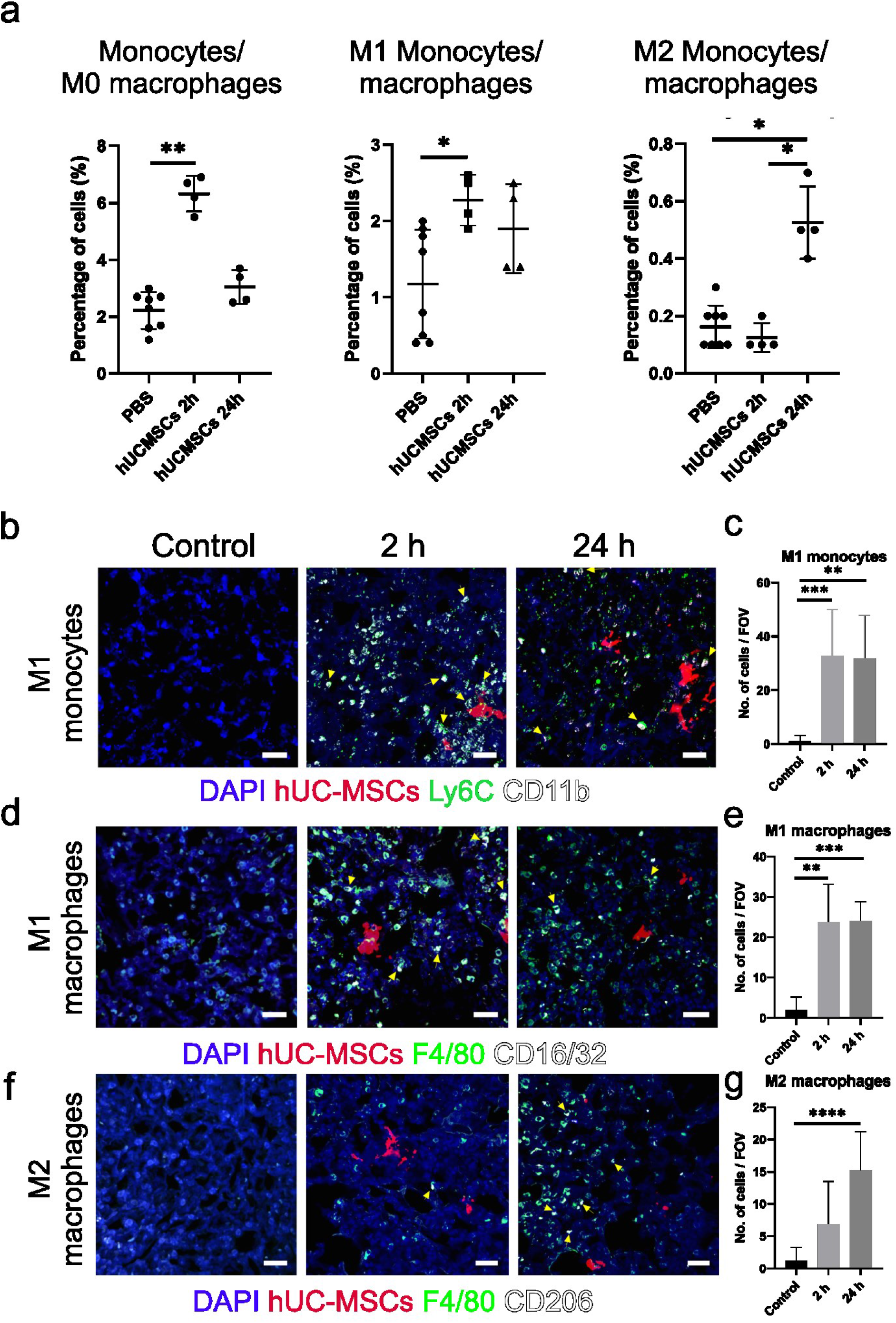
Monocytes and M0 macrophages in the lung show a 2-step polarization response to IV injection of hUC-MSCs. a) Flow cytometry showed that the CD11b^hi^ MHC II^+/-^ CD64^+/-^ monocyte and M0 macrophage populations increased at 2 h and presented a classically activated phenotype. At 24 h these cells transitioned toward an alternatively activated phenotype. Kruskal-Wallis test with multiple comparisons. control n=8, hUC-MSC group n=4. P<0.05 *. b) (CD11b [white] + Ly6C [green]) classically activated monocytes distribution in the lungs (yellow arrows). Scale bar = 30 μm. c) Immunofluorescence quantification of LY6C + CD11b M1 monocytes. Kruskal-Wallis test with multiple comparisons. n=9. P<0.005 **; P<0.0005 ***. d) M1 macrophages (F4/80 [green] + CD16/32 [white]) biodistribution in the lung after cell therapy (yellow arrows). Scale bar = 30 μm. e) Immunofluorescence quantification of F4/80 + CD16/32 M1 macrophages. Kruskal-Wallis test with multiple comparisons. n=9. P<0.005 **; P<0.0005 ***. f) (F4/80 [green] + CD206 [white]) M2 macrophages distribution (yellow arrowheads). Scale bar = 30 μm. g) Immunofluorescence quantification of F4/80 + CD206 M2 macrophages Kruskal-Wallis test with multiple comparisons. n=9. P<0.00005 ****.

To study immune cell polarization within lung tissue sections, we performed co-staining for CD11b and LY6C (M1 monocytes). There was an even distribution of M1 monocytes throughout the tissue without a tendency for them to accumulate around the hUC-MSCs at both time points (Figure 6b). Quantification of M1 monocytes showed an infiltration of these cells into the lung at 2 h which was sustained at 24 h (Figure 6c). M2 monocytes were not investigated by immunofluorescence but it has previously been shown that there is an increase in this cell type at 24 h post cell injection (26).

Co-staining for the F4/80 and CD16/32 markers revealed that M1 macrophages distributed homogeneously throughout the tissue (Figure 6d). Quantification showed that M1 macrophages infiltrate the lung 2 h after cell injection with the level of these cells remaining high at 24 (Figure 6e). To identify M2 macrophages, we used the F4/80 and CD206 markers. Double-labelled cells were observed homogeneously distributed within the tissue at 24 h (Figure 6f) in agreement with the quantification which showed that M2 macrophages increase only after 24 h (Figure 6e).

Taken together, the monocyte/M0 macrophage population was increased significantly 2 h post hUC-MSC injection. At this time point, an inflammatory response was observed as both monocytes and macrophages differentiated toward an M1 phenotype. At 24 h, although the levels of M1 cells remained high, as shown by immunofluorescence, a resolution of inflammation phase was observed as some of the monocytes and macrophages acquired an M2 phenotype.

### 3.6 Effect of hUC-MSCs on the proportion and polarization of lung macrophage subpopulations

As shown in *Figure 6*, IV administration of hUC-MSC increased the overall macrophage levels in the lung. Their rapid increase within a 2 h period suggests that these cells are infiltrating macrophages from bone-marrow and spleen (55). However, it is not clear whether the hUC-MSCs also have any effect on the resident macrophage populations in the lung, which comprise interstitial and alveolar macrophages. To address this, we used flow cytometry to investigate the effect of hUC-MSCs on Cd11b-CD11c^hi^ CD64^+^ alveolar and MHC II^+^ CD11b^+^ CD64^+^ CD24^-^ interstitial macrophages as well as their polarization status.

The analysis showed that although not significant, a trend towards lower cell number was observed at 2 h. The resident alveolar macrophage levels, with just a 0.4x increase at 24 h, remained relatively stable after hUC-MSC infusion (*Figure 7a*, left). With a 0.5x drop in M1 macrophages, these cells showed a similar trend to the overall population. The MSCs did not trigger a pro-inflammatory response in these cells (*Figure 7a*, middle). M2 alveolar macrophages were below the level of detection at 2 h, but the small 0.16x increase at 24 h does not suggest that alveolar macrophages change their phenotype toward an M2 activation state (*Figure 7a*, right). The interstitial macrophage population remained unchanged 2 h post cell injection but increased by 2.8x 24 h later. At this timepoint, polarization towards both an M1 and M2 phenotypes was observed, with increases of 2.3 and 4.2x, respectively (Figure 7b).

**Figure 7.**
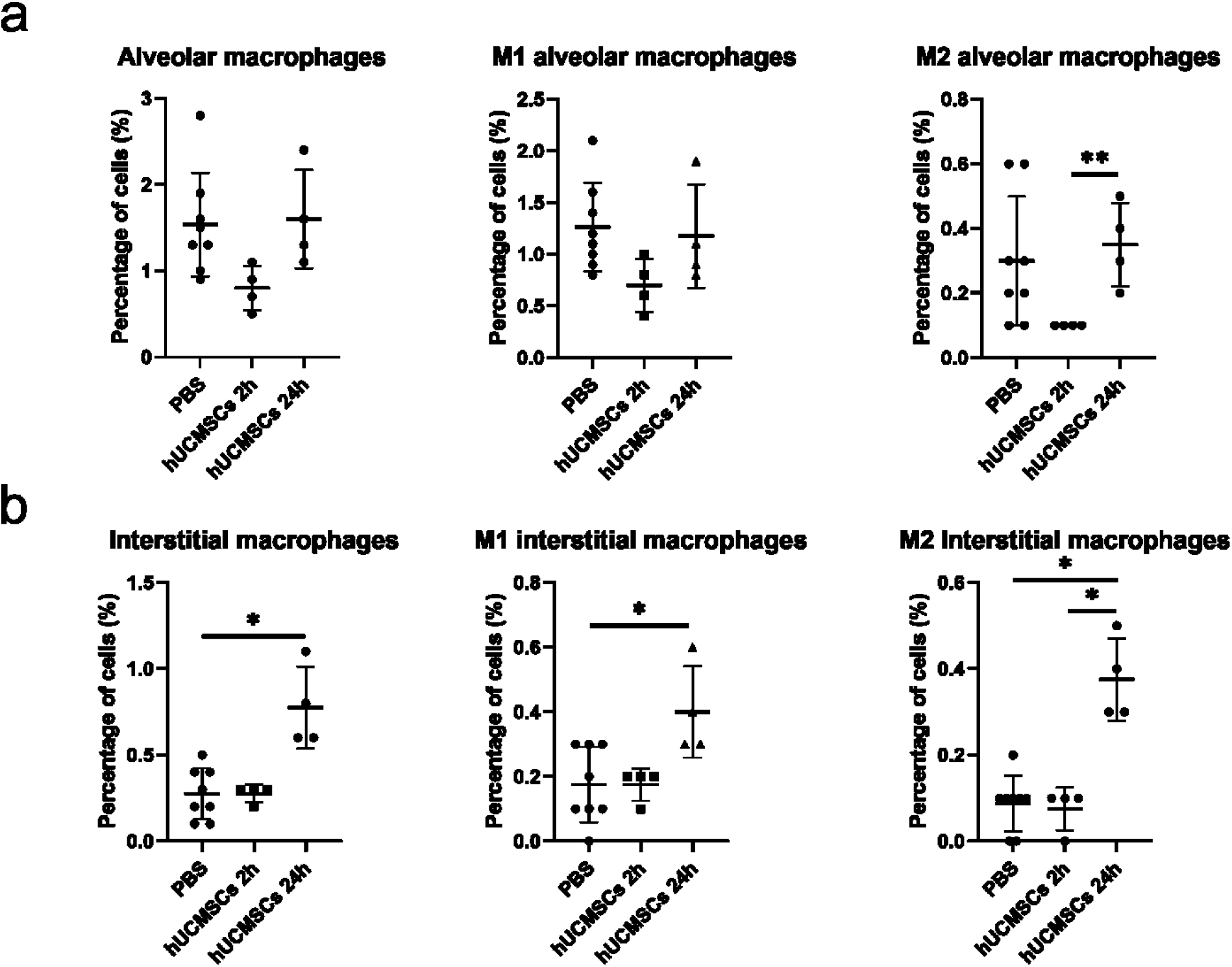
*Macrophage subsets and their polarization after IV administration of hUC-MScs. a) Flow cytometric analysis of* Cd11b^-^ CD11c^hi^ CD64^+^ *alveolar macrophages and their polarisation in the lung b*) MHC II^+^ CD11b^+^ CD64^+^ CD24^-^ *interstitial macrophages were also analysis for changes in their proportion and polarisation by flow cytometry. Kruskal-Wallis test with multiple comparisons. control n=8, hUC-MSC group n=4. P<0.05 * P<0.005 \*\**.

**Figure 8.**
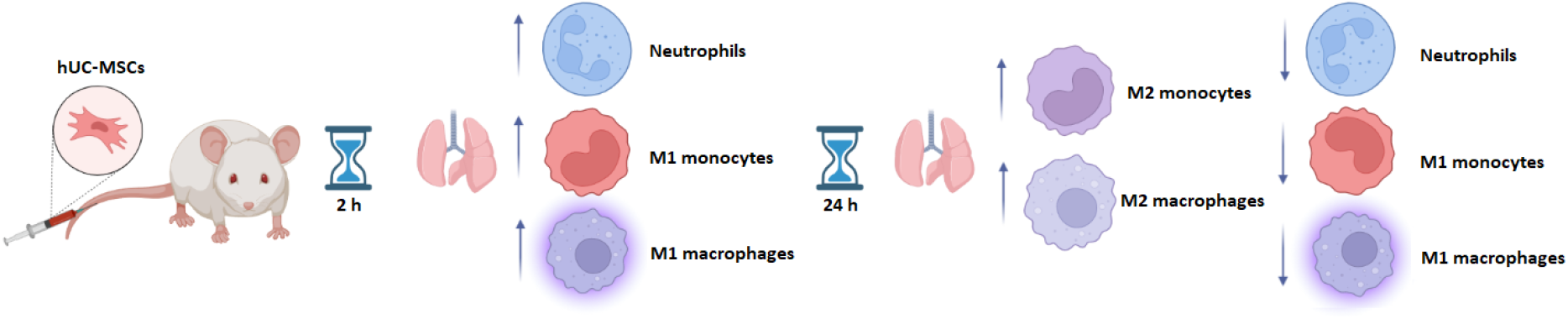
Summary of the effect of IV delivered hUC-MSCs on innate immune cells.

IV administration does not seem to have an effect on alveolar macrophages in the lung. On the contrary, the levels of interstitial macrophages are significantly increased and show an initial inflammatory phase that shifts toward a regulatory phase at 24 h, once most of the infused cells have been cleared.

## 4. Discussion

In this study we observed changes in the proportion of different innate immune cell subtypes after the IV delivery of hUC-MSCs into immunocompetent healthy mice. The key finding was that within 2 h of MSC administration, a pro-inflammatory response was observed in the lung, which appeared to become anti-inflammatory by 24 h. Moreover, histological data, although not directly comparable, corroborated these findings and expanded on the tissue biodistribution of the myeloid cells in relation to the MSCs.

Overall, hUC-MSCs triggered an innate immune response shortly after being infused intravenously. The increase in inflammatory cell types agrees with findings that have shown that MSCs delivered IV induce an inflammatory response in the lung, as evidence by macrophage infiltration (28). Moreover, the localisation of macrophages in close proximity to the MSCs at 2 h post-hUC-MSC administration suggests that the clearance of the exogenous cells from the lungs might involve efferocytosis by phagocytes (56).

After 24 h, the levels of inflammatory cells lowered and a shift of monocytes and macrophages toward an anti-inflammatory phenotype was observed, suggesting the onset of an immune-regulatory phenotype. The production of suppressive myeloid cells after close interaction with MSCs has been described as one of the mechanisms by which the therapeutic cells exert positive disease outcomes (57).

Identifying myeloid cell subtypes and their polarization state in the lung is a complex task. Several myeloid populations express similar and overlapping markers. Moreover, researchers have used inconsistent antibody panels resulting in a lack of strictly defined identifiers for specific cell subsets (58). We followed a well-established protocol (42) and modified it in accordance with our experimental goals, offering a novel approach to studying the changes in the proportion of different myeloid cells response to hUC-MSC delivery in *vivo*.

We and others have shown that hUC-MSC administered IV lead to the cells accumulating in the lungs and are cleared within 24 h (1,3,59,60). Given that the hUC-MSCs are short-lived, their mechanisms of action in the resolution of disease are still unclear. We showed that interactions between transplanted MSCs with phagocytic myeloid cells occur shortly after administration. In agreement with our data, others have shown direct interactions of MSCs with host platelets and neutrophils *in vivo* (61), and that MSCs colocalize with macrophage and granulocyte markers via immunofluorescence *ex vivo* (38), suggesting that MSCs might affect the innate immune system through cell-to-cell interactions (62). Phagocytosis of exogenous MSCs by innate immune cells has been demonstrated *in vivo* (26,63) and it has been found to trigger monocytes to adopt an anti-inflammatory phenotype (26).

Ourselves and other have shown that following IV administration, most MSCs die within 24 h (1,2). It has been suggested that this rapid cell death might be required for the MSCs to exert their benefits (56). After IV injection, MSCs undergo apoptosis in the lung. Their subsequent efferocytosis by macrophages was suggested to be the mechanism behind the reduction in the severity of allergic asthma (64). Moreover, clinical data from patients with graft-versus-host disease (GvHD) who have been administered MSCs intravenously, as well as preclinical murine models showed that the host’s cytotoxic cells actively induce the exogenous MSCs to undergo apoptosis. This results in a recipient-induced immunomodulation which is required for improved outcomes (65). In keeping with the finding that viable MSCs are not required to ameliorate injury, heat-inactivated MSCs were able to maintain their immunomodulatory capacity and reduce sepsis in mice (66,67).

Here, we used xenogeneic cells, but studies that have used syngeneic (28) and allogeneic (68) MSCs have also observed an inflammatory immune reaction after MSC delivery, suggesting that the response is not due to the cells being from another species, but rather a response to the MSCs being present in an atypical location, which initiates a clearance mechanism (28). In line with this, the immune system reacts to cells that are not normally in contact with the bloodstream (69). The direct interaction between the MSCs and the blood immediately after infusion might trigger an instant blood-mediated inflammatory reaction (IBMIR), that would not be expected when administering cells that are normally present in the blood circulation, such as leukocytes (70). IBMIR causes platelet-, coagulation-, and complement activation, and results in the MSCs being destroyed quickly, inducing the innate immune system to eliminate them (71).

Regarding lung resident macrophages, our data showed that the hUC-MSCs did not induce changes in the proportion or phenotype of alveolar macrophages. In contrast, others have shown that MSCs induce a slight increase in alveolar macrophages following IV administration (64). Moreover, alveolar macrophages were observed to efferocytose the exogenous MSCs, which polarised them towards an M2 phenotype (64,72,73). This might be explained by the fact that healthy mice were used in the current study, while the three studies cited above used a mouse model of allergic asthma. Alveolar macrophages are the first immune effector cells at the air-lung interphase (74), meaning that the induction of asthma might have activated, primed and mobilized the alveolar macrophages prior to MSC therapy (75). Thus, it is difficult to attribute the observed effects solely to the delivery of MSCs.

As discussed above, the disease context can influence MSC behaviour (76,77). MSCs can either promote or suppress the immune response as shown by *in vitro* culture of MSCs exposed to different clinical bronchoalveolar lavage (BAL) samples representing a wide range of lung pathologies (77). In our study, healthy animals were used; if injured animals had been studied, it is possible that the results might have differed. Thus, the understanding of the effect of IV delivery of hUC-MSCs on the innate immune system in different disease and inflammatory contexts remains to be elucidated.

Regarding lung interstitial macrophages, we observed an 2.8-fold increase in interstitial macrophages at 24 h, which agrees with Pang, et al. who observed that lung interstitial macrophages play an important role in the clearance of exogenous MSCs (64).

How the immunoregulatory properties of MSCs relates to their beneficial effects in disease and injury models remains unclear. Nevertheless, immunomodulation by innate immune cells mediated by the MSC secretome as well as by direct interaction with viable, apoptotic, inactivated, and fragmented MSCs has been established (56). Importantly, MSCs appear to be able to induce therapeutic effects without long-term engraftment (78).

## 5. Conclusions

We performed a comprehensive flow cytometry and histological analysis of mouse lungs following IV administration of hUC-MSCs to investigate the fate of the hUC-MSCs and their effect on the cells of the innate immune system.

We showed that in healthy, immunocompetent mice, an inflammatory response, dominated by an increase in granulocytes -particularly neutrophils-, and pro-inflammatory monocytes and macrophages in the lungs, occurred 2 h after cell delivery. These cells were frequently observed in proximity to the hUC-MSCs, which may indicate that they participate in their clearance by means of phagocytosis. After 24 h, a resolution of the inflammatory phase was observed as anti-inflammatory monocytes and macrophages became more prevalent in the lung. These processes might be involved in the immunomodulatory response following MSC infusion in models of disease. Further research is necessary to ascertain the exact cause of the immune response to better tailor cell therapies to specific conditions.

## 6. Acknowledgements

Animal procedures in this article were performed in the Centre for Preclinical Imaging (CPI) of the University of Liverpool. The CPI has been funded by a Medical Research Council (MRC) grant (MR/L012707/1). All microscopy data were acquired at the Centre for Cell Imaging (CCI) of the University of Liverpool. The confocal system used in this work was funded by BBSRC grant number BB/R01390X/1. The histology work was performed at the histology facility of the University of Liverpool. The authors gratefully acknowledge these facilities for their support and assistance in this work. We thank Dr Marie Held (CCI) for the expert advice on image analysis and Jennifer Adcott for training and help with the use of the Dragonfly. We kindly thank Dr Helen Wright for providing materials and helpful advice regarding NETs. This work was supported by the European Union’s Horizon 2020 research and innovation programme under the Marie Skłodowska-Curie grant agreement No. 813839.

## 7. Conflicts of interests

The authors declared that they have no conflicts of interest to this work.

## Notes

http://doi.org/10.5281/zenodo.7113094

### Competing Interest Statement

The authors have declared no competing interest.

