## Supplementary information for "Intravenous administration of human umbilical cord mesenchymal stromal cells leads to an inflammatory response in the lung"

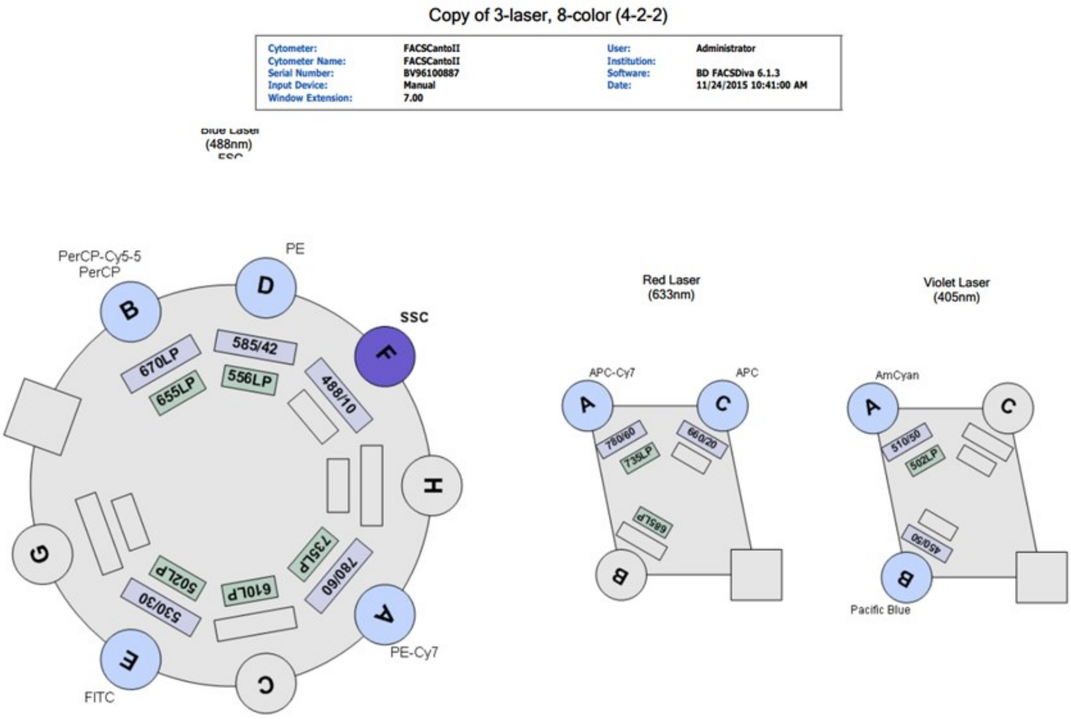


*Supplementary figure 1. BD FACS CANTO II configuration.*

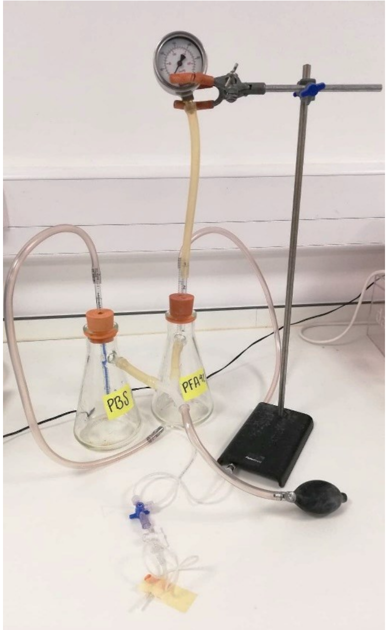


*Supplementary figure 2 Perfusion pump. Two Erlenmeyer flasks, one filled with PBS and the other with 4% PFA, are connected by tubes to their sidearms, which are attached to a manual bulb that helps pressurise the system. One of the bottles has a pressure gauge attached to it to monitor and maintain pressure. Pipettes are inserted into each flask's fluid using rubber stoppers. The liquid from each bottle is pumped into these pipettes and out of the flasks by the pressure flowing through the sidearm, which is controlled by a stopcock that permits fluid to flow from only one flask at a time.*
